# Alignment-free phylogenetic inference via hyperbolic protein language models

**DOI:** 10.64898/2026.05.26.723419

**Authors:** Yongtao Shan, Pan Fang, Yuqi Liu, Yuanfei Pan, Kaijie Liu, Yong He, Xue Liu, Weichen Wu, Guirong Xue, Jian He, Deyin Guo, Jianguo He, Edward C. Holmes, Mang Shi, Zhaorong Li

## Abstract

Conventional phylogenetic methods rely on multiple sequence alignments which are computationally intensive and often fail for highly divergent lineages. Here, we introduce LucaPhylo, an alignment-free framework that infers evolutionary relationships directly from unaligned sequences. Through a cascaded learning strategy LucaPhylo integrates protein language models with hyperbolic geometry, a representation space naturally suited to hierarchical branching, to capture deep evolutionary constraints without explicit homology matching. Using highly divergent RNA virosphere as a test case, LucaPhylo places unaligned sequences into phylogenetic trees with an accuracy comparable to leading alignment-based tree construction tools, while retaining divergent sequences that conventional pipelines frequently discard. It further enables the integration of divergent viral lineages into phylogenetic trees, thereby expanding the evolutionary landscape of RNA viruses. Together, LucaPhylo establishes an AI-driven, alignment-free paradigm for phylogenetic inference and provides a robust computational foundation for resolving deep evolutionary relationships among RNA viruses and other biological systems.

## Introduction

Determining the evolutionary relationships among species is a fundamental goal in biology. Conventional phylogenetic inference relies on multiple sequence alignments (MSAs) followed by model-based tree estimation, typically using computationally intensive approaches such as maximum likelihood (*1*) or Bayesian (*2*) methods. As gene sequence data sets expanded, heuristic search algorithms implemented in software such as PhyML (*3*), RAxML (*4*), and FastTree (*5*) enabled large-scale *de novo* phylogenetic inference, while tree placement tools (e.g., pplacer (*6*), EPA-ng (*7*)) were developed to insert novel sequences into existing reference backbone trees without rebuilding entire phylogenies. Despite these methodological advances, all frameworks ultimately depend on explicit sequence alignment and models of nucleotide or amino acid substitution, and usually rely on heuristic approximations that compromise accuracy at scale (*8, 9*). However, when evolutionary divergence enters the “twilight zone” of low sequence similarity (*10*), alignment artifacts proliferate and substitution-based distances saturate, effectively rendering highly divergent lineages phylogenetically unplaceable (*11*).

Recent advances in biological large language models (LLMs) provide a new paradigm. Pre-trained on massive biological data, foundation models such as the ESM (*12, 13*) and Luca (*14, 15*) families encode deep structural and evolutionary signals directly from raw sequences, enabling the detection of remote homologies (*16*) and the prediction of evolutionary trajectories (*17, 18*). However, the high-dimensional mathematical representations (often termed embeddings) generated by these models are typically compared using Euclidean metrics that inadequately reflect the exponential expansion of phylogenetic tree space (*19*). In contrast, hyperbolic manifolds, characterized by exponentially growing volume, offer a mathematically natural representation space for hierarchical structures (*20*). Although hyperbolic spaces have been explored in phylogenetics, prior efforts relied on shallow encodings and did not leverage the complex evolutionary constraints and hidden structural patterns captured by modern foundation models (*21-23*).

Here, we introduce LucaPhylo, an alignment-free phylogenetic framework that integrates foundational sequence models with hyperbolic geometry to infer phylogenetic relationships directly from unaligned sequences. As a test case for interspecific phylogenetic placement we applied this framework to RNA viruses, a system characterized by high mutation rates leading to extensive sequence divergence (*24*), as well as an ever-expanding sequence database (*25*). By combining a multi-scale representation strategy – from conserved RNA-dependent RNA polymerase (RdRP) domains to full-length polyproteins – our framework enables the accurate phylogenetic placement of both individual sequences and entire clades, including highly divergent taxa systematically discarded in alignment-dependent placement pipelines. LucaPhylo therefore provides a scalable strategy for resolving complex evolutionary placements independent of explicit sequence homology, advancing phylogenetic analyses in virology with the capacity to extend other sets of divergent sequences.

## Results

### The LucaPhylo framework: alignment-free phylogenetics via protein language models

LucaPhylo represents a computational framework that integrates a protein language model (PLM) and hyperbolic space embedding to infer phylogenetic relationships directly from unaligned sequences. It does not require MSAs or conventional phylogenetic tree construction algorithms. Using highly divergent RNA virus sequences as a stringent benchmark, we demonstrate that LucaPhylo can accurately place viral RdRP-containing sequence(s) into established phylogenetic backbones and update these trees with entirely novel viral genus- or family-level lineages (Fig. 1A), in turn enabling the robust inference of evolutionary relationships across both closely and distantly related viral sequences.

**Figure 1.**
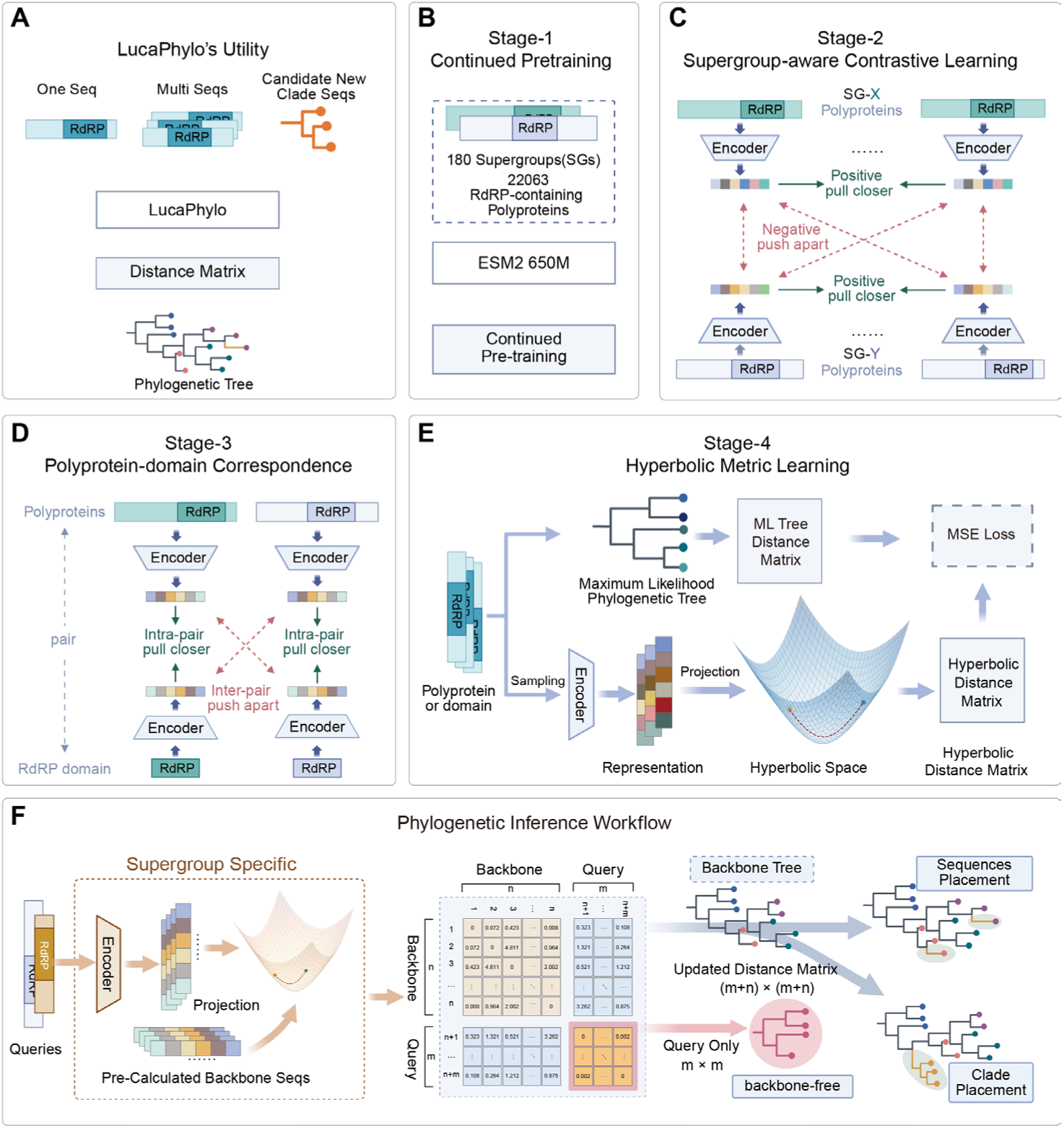
The LucaPhylo framework for alignment-free phylogenetic inference. **(A)** Overview of LucaPhylo’s utility for evolutionary analysis. **(B)** Stage-1: Continued pre-training to adapt the base language model to sequence features specific for RNA viruses. **(C)** Stage-2: Supergroup-aware contrastive learning to encode coarse-grained taxonomic structure. **(D)** Stage-3: Polyprotein-domain correspondence learning, aligning full-length sequences with conserved RdRP domains to fuse multi-scale evolutionary signals. **(E)** Stage-4: Hyperbolic metric learning, where representations are projected into a hyperbolic manifold to capture the hierarchical branching of phylogenetic trees. **(F)** Phylogenetic inference workflow. Unaligned queries and backbone references are encoded into a shared hyperbolic space to construct an updated distance matrix, enabling sequence and clade placement.

To construct an alignment-free framework capable of consistent evolutionary analysis, we aimed to endow the model with the capacity to interpret long polyprotein sequences and autonomously distll critical evolutionary information within a geometrically faithful metric space. To this end, we implemented a four-stage cascaded training strategy that progressively infuses evolutionary structure into the protein language model (Fig. 1A-D). In Stage-1, we performed continued pretraining on RdRP-containing polyproteins using ESM2 650M to adapt the model to sequence features specific to RNA viruses (Fig. 1B). Stage-2 introduced supergroup-aware contrastive fine-tuning. This orientates the high-dimensional feature space along macro-evolutionary directions, thereby encoding coarse-grained taxonomic structure (Fig. 1C). In Stage-3, we jointly optimized full-length polyproteins and RdRP domains to fuse global sequence context with local conserved motifs (Fig. 1D). This multi-scale alignment enables the direct extraction of evolutionary signals from unannotated inputs, effectively bypassing the prerequisites for domain annotation and multiple sequence alignment. Finally, in Stage-4, we projected the learned representations into a hyperbolic phylogenetic manifold under maximum likelihood (ML) tree supervision (Fig. 1E). This transformation enables the direct calculation of evolutionary distances within a geometry that intrinsically accommodates the exponential branching structure of phylogenetic trees.

To ensure generalization across the vast diversity of the RNA virosphere, training data were curated from a large-scale study of RNA virus diversity (*16*), encompassing 21 well-established viral supergroups (i.e. phylum or class level virus groups) as well as 159 newly defined supergroups. The final training set comprised 22,063 RdRP domain sequences and their corresponding polyproteins (Table S1). For each supergroup, ML backbone trees inferred from MSAs were used to supervise training on unaligned sequences.

For inference, LucaPhylo encodes query sequences and backbone references into the same hyperbolic space, from which pairwise evolutionary distances are inferred. These distances are then used to estimate phylogenetic trees or to place novel sequences onto existing phylogenetic backbones (Fig. 1F).

### LucaPhylo discriminates viral supergroups and learns key evolutionary features

We evaluated whether LucaPhylo embeddings capture features of sequence homology across training stages by examining their correspondence to viral supergroups. The Stage-1 model successfully adapted the base model to viral sequences, as indicated by lower perplexity and loss on held-out data (Data set-1, Fig. S1A), thereby providing a stable initialization (Fig. S2).

The Stage-2 model, optimized via supergroup-aware contrastive learning, exhibited coarse-grained taxonomic signatures. For the 21 ICTV-recognized viral supergroups (Data set-2, Fig. S1A; Table S1), Stage-2 effectively captured supergroup-specific signatures. This clustering concordance with ground-truth taxonomy was quantified by an Adjusted Rand Index (ARI) of 0.8881, substantially outperforming the ESM2 650M and 3B(*12*) baselines (Fig. 2A), with clear supergroup separation in low-dimensional projections (Fig. 2B). When extended to a more globally representative and complex data set of 88 supergroups, including 67 newly discovered supergroups (Data set-1, Fig. S1A), the model maintained robust performance (0.8087 ARI; Fig. 2A), demonstrating its scalability across a broader spectrum of viral diversity. These results were further validated by supergroup-based classification analyses using Stage-2 embeddings. Using either cosine distance-based methods or a simple linear classification network (Fig. 2C), Stage-2 embeddings achieved > 91% accuracy on Data set-1 (comprising 88 supergroups; see Table S2 for detailed performance metrics), markedly higher than the baseline models (21.07%–87.39%; Fig. 2D). Notably, this representation generalized to highly divergent viruses lacking detectable sequence homology, indicating that it captures semantic constraints beyond sequence similarity. Specifically, Stage-2 achieved 82.25% accuracy on Data set-3, comprising 11,381 polyproteins with no BLAST hits to the training set, compared with 61.66% for the baseline (Fig. 2E; detailed performance metrics are shown in Table S2). We further calculated the intra- and inter-supergroup cosine distances within Data set-1, revealing that a threshold defined by the maximal mean intra-supergroup distance could reliably separate all supergroups (Fig. S3A, B; detailed performance metrics in Table S3). These results underscore that by learning from representative sequences the model successfully captured the distinctive evolutionary signatures that define each supergroup.

**Figure 2.**
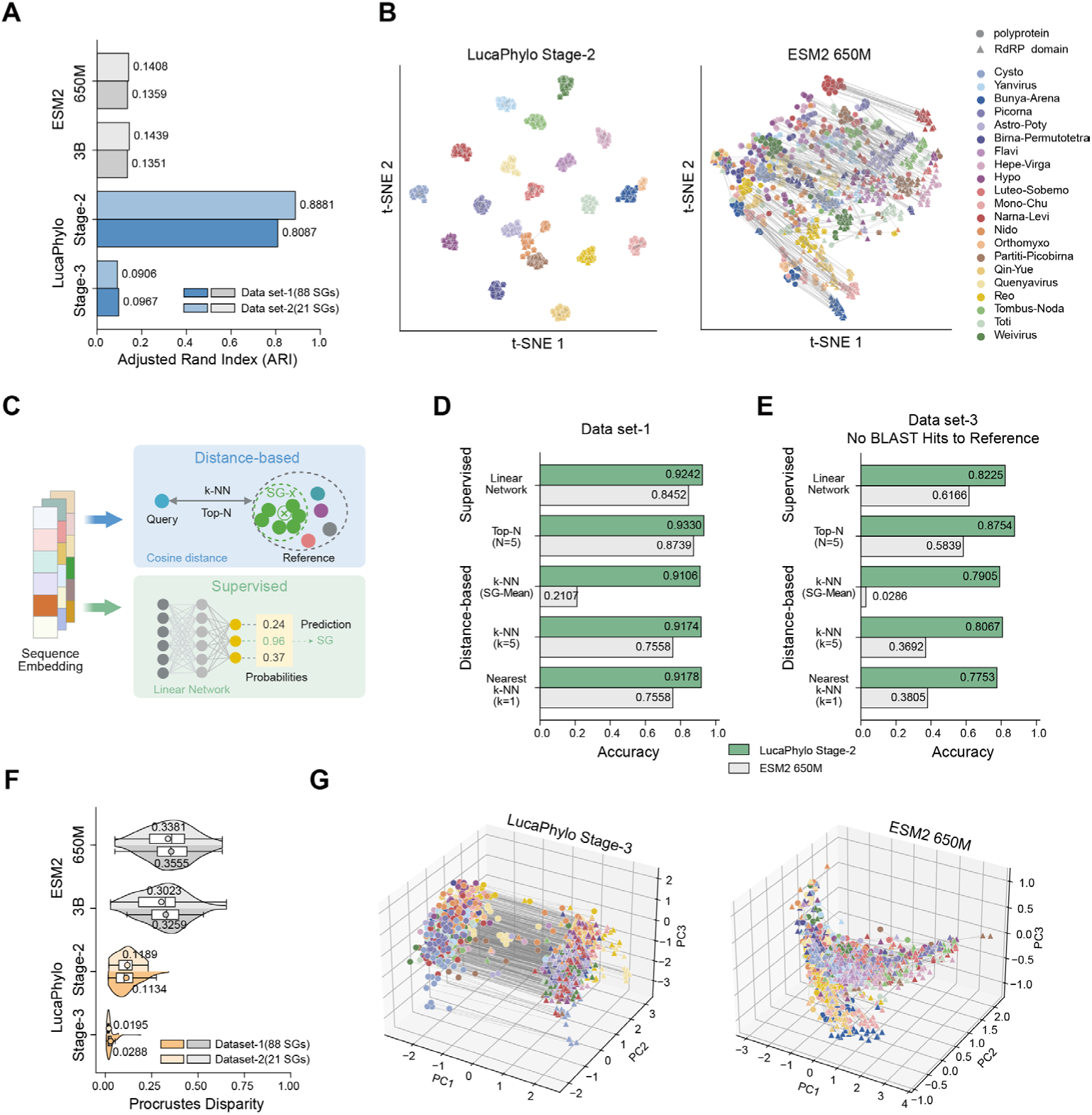
LucaPhylo captures distinct taxonomic signatures and aligns multi-scale evolutionary representations. **(A)** Adjusted Rand Index (ARI) evaluating the clustering concordance of sequence embeddings with ground-truth viral taxonomy. **(B)** Two-dimensional t-SNE projections of sequence embeddings. Colors denote distinct RNA virus supergroups. **(C)** Schematic of supergroup discrimination methodologies, employing zero-shot embedding-based classifiers (cosine distance) and a trained linear classification network. **(D, E)** Classification accuracy of Stage-2 versus the baseline on (D) Data set-1 (representative global viral diversity) and (E) Data set-3 (highly divergent sequences lacking detectable sequence similarity to the training set). **(F)** Distributions of Procrustes distances between matched full-length polyproteins and RdRP domains. Dots and corresponding numerical labels denote the mean values for each distribution. Lower distances indicate stronger structural alignment across scales. **(G)** Three-dimensional PCA visualization of matched polyprotein-domain pairs. Circles and triangles denote polyproteins and corresponding RdRP domains, respectively.

While Stage-2 establishes a robust coarse-grained taxonomic framework, Stage-3 captures fine-grained evolutionary signal by aligning full-length polyproteins with their conserved functional domains (RdRP in this study). This alignment strategy was validated by low Procrustes distances (Fig. 2F) and consistent geometric transformations between matched polyprotein–domain pairs (Fig. 2G), enabling the extraction of domain-level evolutionary signals directly from unannotated polyproteins. The integrated Stage-2 and Stage-3 architecture therefore yielded a multi-scale representation that supports both broad placement of unclassified viral lineages and fine-grained resolution within viral supergroups.

### From embeddings to evolution: learning evolutionary relationships and attention-based insights

We next evaluated the model’s capacity to infer evolutionary relationships without explicit training on specific phylogenetic topologies (i.e., zero-shot inference). Specifically, we investigated whether the latent geometric distances between unaligned sequence embeddings could capture evolutionary divergence, using patristic distances from ML trees as the ground-truth reference. Performance was evaluated against base ESM2 models across two dimensions: local phylogenetic affinity (Jaccard index) and global evolutionary distance concordance (Pearson’s correlation, *r*) (Fig. 3A). In this intra-supergroup inference task, Stage-3 embeddings outperformed all baselines in recovering ground-truth phylogenetic structure across 70 RNA virus supergroups (Data set-4, Fig. S1B; Fig. 3B; detailed performance metrics in Table S4). Crucially, this superiority remained robust across diverse input formats, including MSAs, RdRP domains, and full-length polyproteins (Fig. 3C, D, and Table S5), supporting the model’s alignment-independent and representation-driven inference capacity.

**Figure 3.**
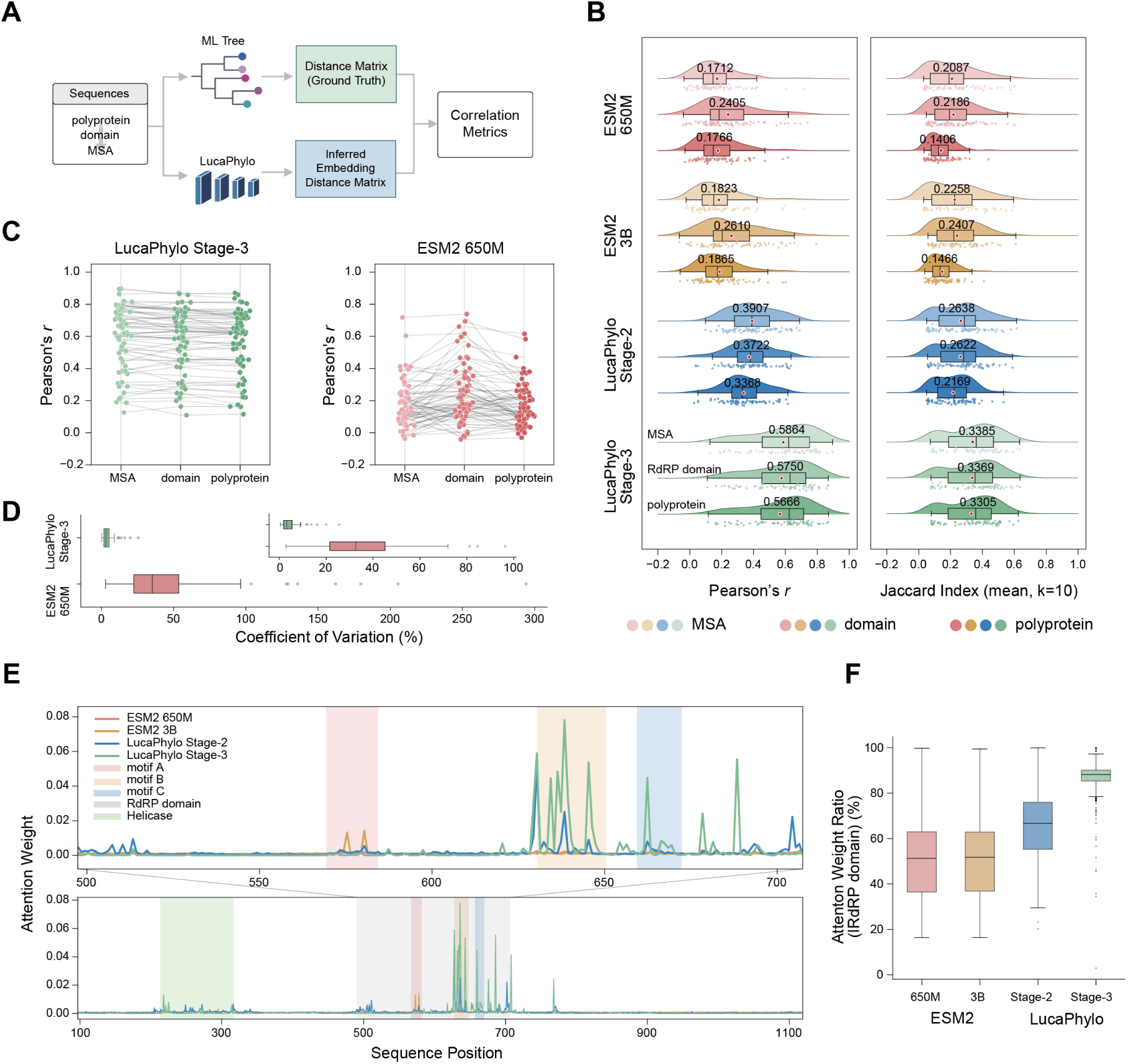
Zero-shot inference of evolutionary relationships and attention-based interpretability. **(A)** Evaluation framework for zero-shot alignment-free phylogenetic inference. **(B)** Performance of intra-supergroup zero-shot inference across Data set-4. **(C, D)** Evolutionary inference performance across diverse input modalities. (C) Distribution of correlation metrics across different models and input types. (D) Coefficient of variation (CV) of Pearson correlation scores. **(E)** Sequence-level attention analysis for a representative viral polyprotein. The plotted values represent the average attention weight received by each residue from all other positions across the sequence. The Stage-3 model (green line) concentrates attention on evolutionarily conserved functional constraints, notably the RdRP domain and auxiliary domains such as the helicase. **(F)** Proportion of total attention weight allocated to the RdRP domain across Data set-2 polyproteins. A higher proportion indicates that the model focuses more on the conserved RdRP domain region.

To explore the representational basis of LucaPhylo’s evolutionary inference, we analyzed its attention mechanism, which maps the internal statistical dependencies between residues during model inference. This revealed that the model’s computational focus was strongly correlated with evolutionarily informative regions. Specifically, the Stage-3 model focuses on conserved functional domains, particularly the RdRP and its three canonical sequence motifs (Fig. 3E). Indeed, 89.55% of the total attention weight in Data set-2 polyproteins was concentrated on the RdRP domain, representing a 62% increase relative to the unoptimized base model (Fig. 3F, and Table S6). Additional attention distributed across auxiliary domains, such as the helicase (Fig. 3E), indicates that LucaPhylo successfully integrates polyprotein-level signals across gene regions. Analyses of the seed aligned Pfam sequences further revealed the positional consistency of high-attention sites across protein families (Fig. S4), suggesting that the model anchors phylogenetic inference to deep, functionally conserved sequence motifs.

### Alignment-free phylogenetic inference beyond sequence homology

To assess whether LucaPhylo can resolve complex phylogenetic placements without explicit sequence alignment, we benchmarked our framework by fine-tuning the Stage-3 base model (via Stage-4 hyperbolic adaptation) to place individual virus sequences or divergent clades into curated reference backbones (Fig. 4A; Fig. S1C, S5A). We then evaluated performance across a two-tiered benchmarking framework: internal similarity-stratified tests and external tool comparisons. Performance was assessed using Pearson’s correlation for evolutionary distance concordance and the Jaccard Generalized Robinson-Foulds (JRF) metric for topological divergence.

**Figure 4.**
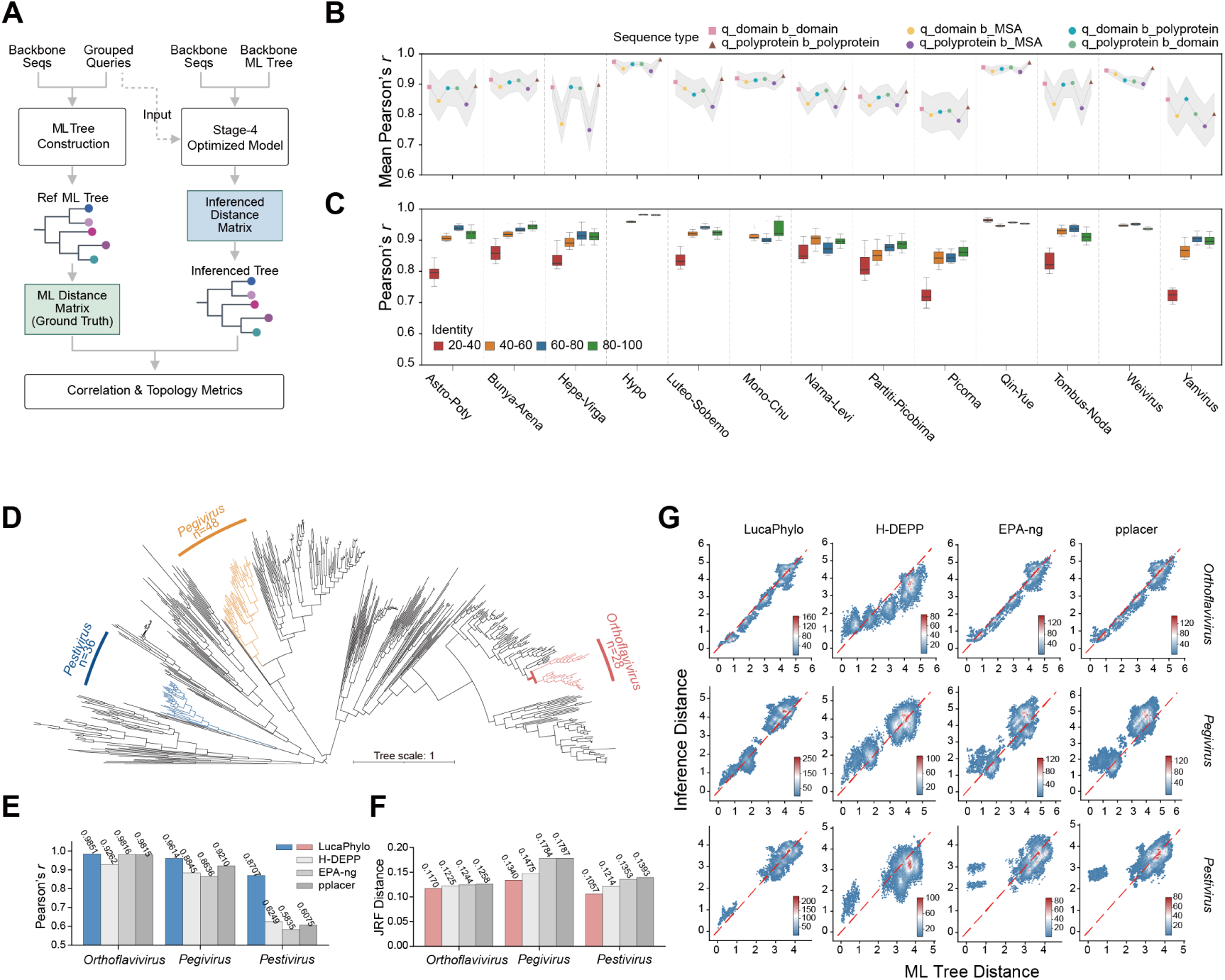
Robustness of LucaPhylo in sequences and clade-level phylogenetic placement. **(A)** Schematic of the evaluation framework for similarity-stratified phylogenetic placement benchmarks. **(B-C)** Correlation performance of the placement of individual sequences by LucaPhylo, with the query set size fixed at 20. (B) Performance across different input modalities. Colors indicate distinct combinations of query (q) and backbone (b) sequence types (e.g., q_domain and b_domain indicates that both are RdRP domains), with shaded regions representing the standard deviation. Data are aggregated across all sequence similarity levels. (C) Performance across varying sequence identities. Analyses were performed using domain sequences for both query and backbone. **(D)** Reference phylogeny of the SG005 (Flavi) supergroup used for clade-placement benchmarks. Colored branches denote the genus-level query sets. **(E, F)** Benchmarking of LucaPhylo against H-DEPP, EPA-ng, and pplacer for the three lineages shown in d. Metrics include (E) Pearson’s r of evolutionary distances and (F) normalized Jaccard Generalized Robinson-Foulds (JRF) metrics. (**G**) Density scatter plots comparing LucaPhylo-inferred evolutionary distances against ground-truth ML patristic distances for the three lineages.

In individual sequence-level placement tests, we first assessed LucaPhylo’s robustness across different input modalities. Notably, LucaPhylo’s alignment-free placements using unaligned polyproteins or conserved domains consistently outperformed its own inferences derived from trimmed MSA backbones (Fig. 4B; Fig. S5B; detailed performance metrics in Table S7), indicating that phylogenetic signal encoded in the learned embedding space is superior to the explicit identification of homology followed by phylogenetic reconstruction. In addition, across controlled gradients of sequence divergence, LucaPhylo demonstrated robust fidelity to ground-truth ML trees, with a performance that was positively correlated with query–backbone similarity (Fig. 4C; Fig. S5C, D; detailed performance metrics in Table S8) and which remained stable across increasing query sizes (Fig. S5E). When benchmarked against established tools (pplacer (*6*), EPA-ng (*7*) and H-DEPP (*21*)) on curated trees for the Flavi (SG005) and Mono-Chu (SG009) supergroups, LucaPhylo outperformed H-DEPP and achieved parity with the other two ML-based methods. Crucially, while existing tools suffered extensive sequence dropout due to alignment failures, LucaPhylo successfully placed all queries, highlighting its advantage in handling sequences that defy conventional alignment (Fig. S6A-D, and Table S9). In addition, computational profiling indicates that LucaPhylo maintains tractable execution times relative to conventional alignment-based pipelines, and enables practical sequence inference across varying query sizes in standard multi-thread CPU environments. For example, the placement of 1,000 unaligned queries onto an 875-sequence backbone took approximately 20 minutes using 32 threads (Fig. S7, and Tables S10, S11).

Beyond individual-sequence(s) placement, LucaPhylo accurately inserted entire divergent lineages, ranging from genera to families, into existing phylogenetic backbones. In clade-level benchmarks, LucaPhylo accurately resolved distinct genus-level lineages (*Pestivirus*, *Pegivirus*, *Orthoflavivirus*) within the curated supergroup SG005 (Flavi) backbone, achieving high correlations (*r* = 0.8707–0.9851) and low topological divergence (JRF = 0.1057–0.1340; Fig. 4D–G). Similarly, it successfully placed distant family-level clades (*Bornaviridae*, *Chuviridae*-subset, *Filoviridae*, *Paramyxoviridae*) into the curated SG009 (Mono-chu) backbone (Fig. S6E–G). Most notably, LucaPhylo uniquely resolved a highly divergent *Mymonaviridae*-like clade that was largely discarded in alignment-based pipelines (*r* = 0.9712; JRF = 0.1290; Fig. S6H, and Table S12).

Together, these results demonstrate that by leveraging learned representations within a hyperbolic space, LucaPhylo accurately infers phylogenetic relationships directly from diverse unaligned inputs, including conserved domains and full-length polyproteins. This enables both individual-sequence and clade-level phylogenetic placements, thereby providing a computational foundation for resolving the deep phylogenetic relationships among viruses that is inaccessible to conventional alignment-dependent placement methods.

## Discussion

Reliable phylogenetic inference has long been constrained by the inability of multiple sequence alignments to perform with accuracy on highly divergent sequences, with failure commonplace when sequence similarity enters the “twilight zone” of low sequence identity (*10, 26*). Herein, we demonstrate that the placement of highly divergent taxa can be robustly performed without strict dependence on explicit sequence alignments. By integrating representations from protein language models with the natural mathematical geometry of hyperbolic manifolds, LucaPhylo transitions phylogenetic analysis from alignment-based sequence similarity towards representations grounded in conserved functional regions and underlying evolutionary constraints. This strategy avoids alignment artifacts while providing a scalable solution that complements conventional model-based phylogenetics.

Conceptually, LucaPhylo reframes phylogenetic inference as a representation learning problem. Traditional phylogenetic analysis typically involves a multi-step pipeline—typically including homologous sequence identification, multiple sequence alignment, and model-based tree inference—each requiring substantial computational resources and manual parameterization (*8, 27*). LucaPhylo implicitly recapitulates these steps through a multi-stage training strategy, but performs them within a unified learning framework. LucaPhylo first captures homologous sequence relationships, enabling divergent yet evolutionarily related sequences to be grouped prior to phylogenetic inference. It then identifies conserved functional regions that serve as comparable evolutionary signals, effectively fulfilling the role of alignment without relying on explicit alignment algorithms. Finally, it translates these learned representations into phylogenetically meaningful distances for tree estimation. In this way, LucaPhylo mirrors the conceptual structure of classical phylogenetic workflows, while achieving the same objectives through an alternative, data-driven pathway that reduces reliance on manual preprocessing and computationally intensive alignment procedures (*28*).

Translating continuous embeddings into accurate evolutionary topologies relies on an appropriate geometric space. Traditional Euclidean representations compress hierarchical structures and lead to spatial crowding among deeply divergent taxa (*19*). LucaPhylo mitigates this by embedding sequences within a hyperbolic manifold whose exponentially expanding volume naturally reflects the branching structure of evolutionary trees (*20*). Consequently, projecting evolutionarily constrained features—such as conserved functional domains—into this geometry preserves large-scale tree topology while capturing long-range evolutionary constraints typically lost in shallow composition-based methods (*9, 29, 30*).

The highly divergent sequences that characterise RNA viruses serve as a major stress test for phylogenetic inference. LucaPhylo facilitates the phylogenetic placement of divergent viral taxa by leveraging signals centered on the RdRP, the only universally conserved viral marker (*31-33*). Crucially, the application to RNA viruses illustrates a broadly generalizable paradigm, as LucaPhylo is not taxon-specific and can potentially extend across the tree of life. Replacing the underlying protein language model with nucleotide-supported foundation models (*14*) further expands its applicability to nucleic acid analyses. Although nucleotide sequences encode weaker structural semantics than proteins (*34*), they may still prove valuable for such tasks as the high-throughput phylogenetic anchoring of microbial communities using 16S rRNA or other conserved markers.

Extending LucaPhylo to multi-gene or whole-genome phylogenetic analyses represents an important future direction. However, modeling reticulate evolution driven by horizontal gene transfer and genomic rearrangements remains challenging (*35-37*), and encoding extremely long genomic sequences represents computational bottleneck (*38*). Future architectures may leverage sparse attention mechanisms (*39*) to handle extended genomic sequences, while Mixture-of-Experts (MoE) models (*40, 41*) could enable biologically informed routing of distinct genomic modules into specialized representation spaces. Successful application to different type of organism or genome-scale data sets will also depend on increasingly comprehensive reference phylogenies for supervising the underlying geometric manifold.

We utilized ML trees inferred from empirical data as evolutionary constraints to guide manifold learning. Although LucaPhylo extracts robust evolutionary representations and accurate pairwise distances, such reference trees do not necessarily represent "true" phylogenies. Simulated data sets offer theoretical ground truth, yet current sequence simulators rely on site-independent substitution models that fail to recapitulate the complex epistatic interactions and deep structural constraints observed in natural evolution (42, 43). Protein language models inherently depend on these authentic evolutionary constraints and conserved patterns to capture deep functional signatures; thus, guiding manifold learning with empirical data ensures that the model maps a viable sequence space shaped by natural selection (*17, 30*). Incorporating broader empirical training data and orthogonal biological priors, including structural similarity information (*42*), will further refine the learned manifold.

Foundation models also risk embedding out-of-distribution (OOD) sequences into inappropriate regions of latent space (*43*). LucaPhylo currently mitigates this through alignment-based screening with representation-based classification. However, defining absolute geometric rejection boundaries remains difficult due to sparse sampling of the evolutionary landscape (*44*). As phylogenetic data sets expand, density-based uncertainty estimation (*45*) may enable models to more reliably identify or reject OOD sequences.

In summary, LucaPhylo establishes an alignment-free paradigm for phylogenetic inference, shifting the analytical basis from sequence homology to continuous, representation-driven evolutionary modeling. By integrating biological foundation models with hyperbolic geometry, the LucaPhylo framework successfully resolves the phylogenetic relationships among highly divergent sequences that remain inaccessible to conventional pipelines. As genomic data continues to expand exponentially, LucaPhylo offers a scalable strategy for translating large unannotated sequence data sets into coherent evolutionary structure. More broadly, our results suggest that geometric representations capture deep biological constraints, opening new opportunities for applications ranging from pathogen surveillance to resolving complex evolutionary relationships across diverse biological systems.

## Materials and Methods

### Data acquisition and data set construction

Source data. Training and external data sets were derived from a previously published global RNA virosphere resource (*16*). The primary training data set comprised representative RdRP domain sequences and their corresponding full-length polyproteins from 180 viral supergroups (22,063 domain-polyprotein pairs). The external data set was constructed by excluding the training sequences and removing duplicates, yielding 425,577 domain–polyprotein pairs spanning 167 supergroups (Fig. S1A).

Representation validation data sets. Three evaluation data sets (Data sets 1-3, Fig. S1A) were generated by stratified subsampling of the external data set.

For Data set-1, all supergroups containing ≥15 sequences were included, with those exceeding 30 sequences randomly subsampled to a maximum of 30 representatives.

Data set-2 is a subset of Data set-1 and restricted to the 21 supergroups with established ICTV taxonomy.

Data set-3 was constructed to assess performance on divergent sequences. Polyproteins in the external data set were searched against the training set using BLASTp (E-value < 1e-5). Sequences without significant hits were retained, resulting in 11,381 sequences across 36 supergroups.

Phylogeny associated data sets. To evaluate the model’s capacity for zero-shot phylogenetic inference within established viral supergroups, we constructed Data set-4 (Fig. S1B) derived from the curated training set. Supergroups containing at least 100 sequences were retained to ensure statistical robustness. Since training only incorporated supergroup labels and not intra-supergroup phylogenetic structure, these sequences serve as a valid zero-shot benchmark. For each supergroup, reference ML trees were reconstructed from RdRP domains. Sequence alignments were generated using MAFFT (v7.525, --maxiterate 1000 --localpair) (*46*) and trimmed with trimAl (v1.5, -gt 0.9 -cons 5) (*47*). Phylogenetic trees were then inferred using IQ-TREE (multicore v2.3.6, -m LG --alrt 1000 -B 1000) (*48*).

Detailed sequence and compositional information for these data sets are provided in Data S1.

### Alignment-free placement benchmarks

Alignment-free placement performance was evaluated through a two-tiered benchmarking framework, encompassing both internal validation and external comparative assessments (Fig. S1C).

Similarity-stratified benchmarks were designed to internally quantify LucaPhylo’s placement accuracy across strictly controlled gradients of sequence divergence. Thirteen supergroups with established ICTV taxonomy in the training set were expanded with additional sequences from the external data set to infer higher-resolution reference phylogenies using the standardized ML pipeline described above, with alignment quality manually inspected. Domain sequences from the external data set were aligned to reference domains using DIAMOND blastp (v2.1.11.165, e-value < 1e-5) (*49*) and stratified by sequence similarity (20–40%, 40–60%, 60–80%, and 80–100%). Within each bin, 5, 10, and 20 query sequences were sampled (20 replications per configuration), and corresponding polyprotein were retrieved. Ground-truth trees were reconstructed by combining the query and backbone domain sequences using the same pipeline without additional manual intervention. Supergroups with insufficient representation from the external data set were excluded, and several lacked sequences in the lowest-identity bin.

Complementing this internal profiling, individual- and clade-level benchmarks were constructed to enable direct comparison with alignment-dependent placement tools and to evaluate the model’s robustness to entire lineage placements. High-resolution reference trees were constructed for supergroups SG005 (Flavi) and SG009 (Mono-Chu) by incorporating taxonomy-annotated sequences from previous studies (*50, 51*), including well-supported monophyletic groups. For the individual-level benchmark, 50 query sequences were randomly sampled from the reference trees, spanning a representative range of phylogenetic distances, with the remaining taxa serving as the backbone. For the clade-level evaluation, entire clades were designated as queries. In both cases, ground-truth trees were reconstructed from the combined backbone and query sets using the same ML pipeline.

### Model architecture and training

LucaPhylo was initialized from the pre-trained ESM-2 model (650M parameters; 33 layers, 1,280-dimensional embeddings), a transformer-based protein language model trained on over 60 million sequences (*12*). A four-stage cascaded training strategy was implemented to progressively refine evolutionary representations. All training was conducted on NVIDIA A100 GPUs.

Stage-1: Continued pretraining. To adapt the general-purpose model to viral evolutionary semantics, we performed continued pretraining using polyprotein sequences from the training data set. The model was trained via a standard Masked Language Modeling (MLM) objective, with 15% of the input tokens randomly masked. Training was executed for 25 epochs at a learning rate of 1e-4.

Stage-2: Supergroup-aware contrastive learning. To encode viral supergroup taxonomy into embedding space, we implemented a supervised, in-batch contrastive learning framework. Sequence embeddings were derived via mean pooling of final-layer hidden states. For a global batch of 𝑁 sequence pairs, positive pairs 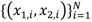 were constructed by independently sampling two distinct sequences belonging to the same viral supergroup, while all other unpaired sequences within the distributed global batch served as negatives. Let 𝐳_1,𝑖_ and 𝐳_2,𝑖_ denote the mean-pooled embeddings for a given pair. The representations were optimized by minimizing the InfoNCE loss (*52*):

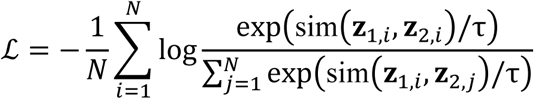

where sim 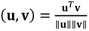 is the cosine similarity, 𝑁 is the global batch size aggregated across all GPUs, and the temperature parameter τ was set to 0.1. The model was optimized for 105 epochs at a learning rate of 1e-5.

Stage-3: Polyprotein–domain correspondence. To establish a correspondence between global sequence context and local conserved motifs while preserving their distinct representational spaces, full-length polyproteins and their corresponding RdRP domains were jointly optimized using a cross-modal contrastive learning framework. For a global batch of 𝑁 matched polyprotein–domain pairs, let 𝐳_𝑝𝑜𝑙𝑦,𝑖_ and 𝐳_𝑑𝑜𝑚,𝑖_ denote their respective mean-pooled embeddings. The primary objective minimizes the InfoNCE loss:

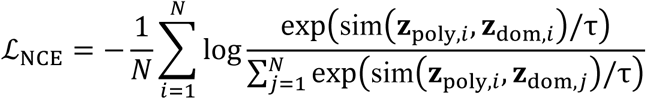

To prevent representation collapse and maintain modality-specific macro-structures, we introduced an orthogonal regularization penalty (ℒ_ortho_) and a centroid separation penalty (ℒ_sep_). ℒ_ortho_enforces embedding diversity within each modality using the Frobenius norm (∥ ⋅ ∥_𝐹_):

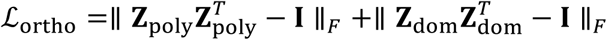

where Z_poly_ and Z_dom_ are the 𝐿_2_-normalized embedding matrices for the batch, and 𝐼 is the identity matrix. ℒ_sep_ penalizes the absolute cosine similarity between the global geometric centers of the two modalities:

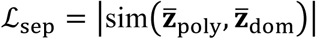

where 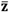 denotes the mean vector across the batch. The overall objective ℒ_total_ = ℒ_NCE_ + λ_ortho_ℒ_ortho_ + λ_sep_ℒ_sep_ was optimized with λ_ortho_ = 0.3, λ_sep_ = 2, 𝑎𝑛𝑑 τ = 0.07. The model was trained for 5 epochs at a learning rate of 5e-5.

Stage 4: Hyperbolic metric learning. To mathematically accommodate the hierarchical branching of phylogenetic trees we projected the learned Euclidean embeddings onto the forward sheet of the Lorentz (hyperboloid) manifold. Specifically, a 𝑑-dimensional Euclidean sequence representation 𝐡 was first mapped into a (𝑑 + 1)-dimensional coordinate space via a learnable linear transformation: 𝐯 = W𝐡 + 𝐛. To rigorously enforce the standard Lorentz manifold constraint, the spatial components were derived directly from this projection (𝐱_1:𝑑_ = 𝐯_1:𝑑_), while the time component (𝑥_0_) was analytically computed as 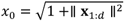. The final hyperbolic representation was thus formed as 𝐱_ℒ_ = [𝑥_0_, 𝐱_1:𝑑_]. The geometric relationship between two representations 𝐱_ℒ_ and 𝐲_ℒ_ on this manifold was quantified using the Minkowski inner product, defined as 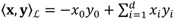.

To accommodate the vastly different scales of evolutionary divergence across taxa we modeled phylogenetic distances as parameterized hyperbolic geodesics. A learnable curvature parameter, 𝑐, was introduced to dynamically scale the global metric space. The scaled hyperbolic distance 𝑑_𝑐_(𝐱, 𝐲) was calculated as:

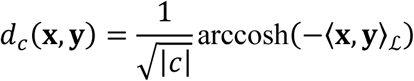

In practice, the negative inner product was set to 1.0 + 1e-6 to prevent gradient singularities. The model was optimized via Mean Squared Error (MSE) regression against ground-truth maximum-likelihood patristic distances. To ensure computational efficiency and preserve pre-trained semantics, we applied Low-Rank Adaptation (LoRA; rank 𝑟 = 16, α = 32) (*53*) to the attention layers while fixing the primary backbone. The AdamW optimizer (learning rate = 5e-5) was employed with a differential learning rate strategy, updating the curvature parameter at 0.1 times the base rate to maintain geometric stability within the hyperbolic space.

### Embedding characterization

To evaluate the quality of the learned representations, sequence embeddings were extracted from the final hidden layer of LucaPhylo (Stage-2 and 3) as well as the baseline models (ESM2-650M and ESM2-3B). High-dimensional representations were projected onto a two-dimensional manifold using t-SNE for visual inspection. Concordance between embedding clusters and established viral taxonomy was quantified using the Adjusted Rand Index (ARI) following k-means clustering, with k set to the number of ground-truth supergroups.

To assess the geometric alignment between polyproteins and their corresponding RdRP domains following Stage-3 training, Procrustes analysis was performed on paired embeddings. We calculated the Procrustes disparity to quantify the structural congruence of these two modalities within the shared latent space. Principal component analysis (PCA) was employed to visualize the transformation vectors connecting these matched pairs, with the aim of assessing whether domain-level evolutionary signals were structurally preserved within the polyprotein embeddings.

### Viral supergroup discrimination

We evaluated the discriminatory capability of LucaPhylo embeddings using both trained linear network and distance-based approaches.

Supervised Classification: A lightweight linear classification network was trained on fixed polyprotein embeddings, which were extracted via mean-pooling of the final transformer layer outputs. The network architecture comprised three hidden layers (dimensions: 1024, 512, and 256), each followed by Layer Normalization, ReLU activation and dropout (probability = 0.2) to mitigate overfitting. Training and validation sets were constructed from the polyprotein training data set using an 8:2 random split. To account for class imbalance across viral supergroups, class-weighted cross-entropy loss was applied with weights inversely proportional to class frequency. The network was optimized using AdamW (learning rate=5e-5) for up to 200 epochs, governed by an early stopping mechanism (patience = 20 epochs) monitoring the validation Macro-F1 score.

Distance-based Classification: We employed zero-shot inference based on cosine similarity to evaluate classification performance without parametric tuning. The embedding space was assessed using the following strategies: (i) Nearest Neighbor (1-NN), which assigns the query to the supergroup of its single closest reference sequence; (ii) k-Nearest Neighbors (k-NN) and Mean Distance, which determine the assignment either via majority voting among the top-k neighbors or by selecting the supergroup with the minimum average distance to the query; and (iii) Top-5 Accuracy, which defines a prediction as successful if the ground-truth supergroup was present among the five nearest reference sequences.

Independent of external references, intrinsic embedding separability was evaluated by computing pairwise cosine distances among all test sequences. Raw distances (𝑑) were transformed into logarithmic similarity scores using 𝑠 = − ln(𝑑 + 𝜀), where 𝜀 was set to 1e-10 to ensure numerical stability. Similarity scores were stratified into intra-supergroup and inter-supergroup categories. For each supergroup, the mean intra-supergroup similarity score was calculated and a global separation threshold was defined as the minimum of these values across all supergroups, representing the lower bound of within-group cohesion.

### Evolutionary concordance analysis

Evolutionary concordance between embedding space and phylogenetic structure was assessed by comparing pairwise cosine distance matrices derived from mean-pooled sequence embeddings with patristic distance matrix obtained from ML trees for each viral supergroup. The correspondence between these matrices was quantified using Pearson correlation coefficients and by calculating the average Jaccard index of *k*-nearest neighbors across varying neighborhood sizes (*k* =5, 10, 20). Robustness to input variations was examined across three distinct sequence types: full-length polyproteins, RdRP domains, and trimmed MSAs. The stability of the phylogenetic signal was quantified using the coefficient of variation (CV) of the Pearson correlation scores across input types.

### Attention-based Interpretability Analysis

To investigate the biological relevance of the learned representations, attention patterns from the final transformer layer of LucaPhylo (Stage-2 and 3) were analyzed and compared with baseline models (ESM2 650M and ESM2 3B). For a given input sequence of length 𝐿, attention weights from all 𝐻 heads in the final layer were first averaged to obtain a unified residue-to-residue attention matrix 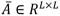. The sequence-level functional importance of the 𝑗-th residue, denoted as 𝑠_𝑗_, was then calculated as the column-wise mean of 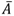, representing the average attention it received from all other positions across the sequence:

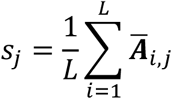

Special tokens located at the sequence termini were strictly excluded from downstream quantitative analyses. Attention profiles were generated for a representative viral polyprotein (YP_009333315.1) and Pfam seed sequences (PF00680) (*54*). Annotated functional domains (RdRP, Helicase) and conserved catalytic motifs (RdRP motifs A, B, C) were mapped onto these profiles to enable comparison of attention allocation across models. Let 𝒟_RdRP_ denote the set of positional indices corresponding to the annotated RdRP domain, and 𝒱 denote the set of valid amino acid indices. The ratio was defined as the cumulative attention score within the RdRP domain normalized by the total attention across the valid polyprotein sequence:

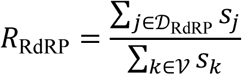

The distributions of these ratios were comprehensively compared across models.

### Tree-specific training and inference

For tree-specific evolutionary distance metric learning, Stage-2 or Stage-3 models were fine-tuned using the hyperbolic optimization strategy described in Stage-4. Representative data sets were constructed using distance-stratified sampling, where training and validation sets (9:1 ratio) were selected at equidistant intervals based on sorted evolutionary distances relative to seed sequences from the reference ML backbone. Models were trained for up to 10 epochs, guided by an early stopping mechanism (patience = 3) monitoring the validation loss.

During inference, query and backbone sequences were projected into the shared hyperbolic space to estimate pairwise geodesic distances. For phylogenetic placement, a matrix augmentation strategy was employed to construct a hybrid distance matrix that integrates the learned representations while strictly preserving the evolutionary topology of the reference scaffold. Let 𝑁_B_ and 𝑁_Q_ denote the number of backbone and query sequences, respectively. The augmented distance matrix 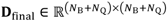 was constructed as a symmetric block matrix:

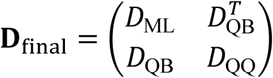

where 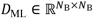 is the ground-truth patristic distance matrix extracted directly from the reference ML tree, anchoring the global topology. The blocks 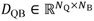 and 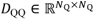 comprise the pairwise hyperbolic geodesic distances between the query and backbone sequences, as well as among the query sequences themselves, dynamically inferred by LucaPhylo.

For query-only reconstruction, distance matrices were derived exclusively from model predictions (i.e., 𝐷_QQ_). Final phylogenetic trees were estimated from the complete distance matrices using FastME (v2.1.6.2, -s -n O) (*55*).

### Placement benchmarking

LucaPhylo was benchmarked against three phylogenetic placement tools: pplacer (v1.1.alpha20) (*6*), EPA-ng (v0.3.8) (*7*), and H-DEPP (v0.1.36) (*21*). As these methods require pre-aligned queries, all query sequences were aligned to backbone MSAs using UPP (v4.5.6) (*56*) with default parameters. pplacer was executed using default parameters. EPA-ng was run using the LG substitution model, matching the model used for inference of the backbone ML tree. H-DEPP was trained with an embedding dimension of 1280 for 10,000 epochs, followed by tree estimation using FastME under the same pipeline.

Performance was assessed using evolutionary distance correlations (Pearson’s *r*) and topological divergence (the Jaccard Generalized Robinson-Foulds metric, JRF) (*57*). Pearson correlations were calculated between inferred and ground-truth ML patristic distances, strictly restricted to query-backbone and query-query pairs. Topological divergence was quantified using the normalized JRF calculated by TreeDist (v2.11.1) (*58*) to provide a robust measure of tree concordance. Notably, JRF metrics were computed only for trials in which all query sequences were successfully placed.

### Computational efficiency evaluation

The computational efficiency of LucaPhylo was evaluated through a two-tiered approach: a benchmark against established alignment-based placement tools and an internal assessment across varying configurations. Both evaluations simulated realistic analytical workflows by placing novel query sequences onto pre-computed phylogenetic reference backbones.

Conventional tools suffered extensive sequence dropout due to alignment failures when processing divergent queries from the external data set. The comparative benchmark was restricted to the curated Flavi (SG005) reference backbone data set (backbone comprising 536 sequences) described in sequence-level placement, only utilizing query subsets capable of successful alignment to the backbone MSA. From this subset, query cohorts of 10, 20, 30, and 40 sequences were randomly sampled (five independent replicates per cohort size). Baseline methods (EPA-ng, pplacer, and H-DEPP) required pre-aligned queries (mean length: 206.4 amino acids), whereas LucaPhylo directly processed raw, unaligned sequences (mean length: 259.4 amino acids). All benchmarks were executed in a standardized 32-thread CPU environment. To ensure equitable evaluation, "total time" was defined as the complete process: comprising explicit pre-alignment and tree insertion for conventional tools, versus direct sequence encoding, distance calculation, and tree estimation for LucaPhylo.

To further evaluate LucaPhylo’s computational performance across varying query sizes, tests were conducted on two empirical backbones comprising 486 and 875 sequences, respectively. Query cohorts of 50, 100, 300, 500, and 1,000 unaligned sequences were randomly sampled from the external data set (10 replicates per cohort). Executions were performed across a single NVIDIA A100 GPU, as well as 16-thread and 32-thread CPU environments. Two metrics were recorded: inference time (query encoding and hyperbolic distance matrix calculation) and total time (incorporating the subsequent FastME tree estimation).

## Supporting information

Figures S1 to S7; Legends for Tables S1 to S12; Legends for Datasets S1.

## Acknowledgments

We thank S-QM from Sun Yat-sen University for valuable discussions regarding the downstream tasks. We thank Y-HW, YS for their on-site technical support and assistance with computational infrastructure maintenance. We also acknowledge the Yunqi Academy of Engineering (Hangzhou) for providing the computational resources that supported this work.

## Author Contributions

Conceptualization: MS, Z-RL; Methodology: Y-TS, PF; Investigation: Y-TS, PF, Y-QL, K-JL; Resources-Computational: YH, G-RX, MS, Z-RL; Project administration: Y-FP, MS, Z-RL; Supervision: J-GH, ECH, MS, Z-R; Writing – original draft: Y-TS, MS, Z-RL; Writing – review & editing: All authors

## Data availability

All data, code, and materials used in this analysis are publicly available to researchers for reproducing or extending the findings. The source code for the LucaPhylo framework, including the multi-stage training and alignment-free phylogenetic inference, along with instructions for obtaining trained weights, is accessible at https://github.com/NickShannn/LucaPhylo. The curated data sets are provided in Data S1. The training data, external data, and all associated model weights—including the foundational weights from the first three representation learning stages, as well as the pre-trained, supergroup-specific hyperbolic model weights ready for direct RNA virus tree inference—are accessible on Zenodo (DOI: 10.5281/zenodo.19401368).

