## Supplementary material for "Alignment-free phylogenetic inference via hyperbolic protein language models": Figures S1 to S7; Legends for Tables S1 to S12; Legends for Datasets S1.

**This PDF file includes:**

Figures S1 to S7

Legends for Tables S1 to S12

Legends for Datasets S1

**Other supporting materials for this manuscript include the following:**

Tables S1 to S12

Datasets S1

### Figures

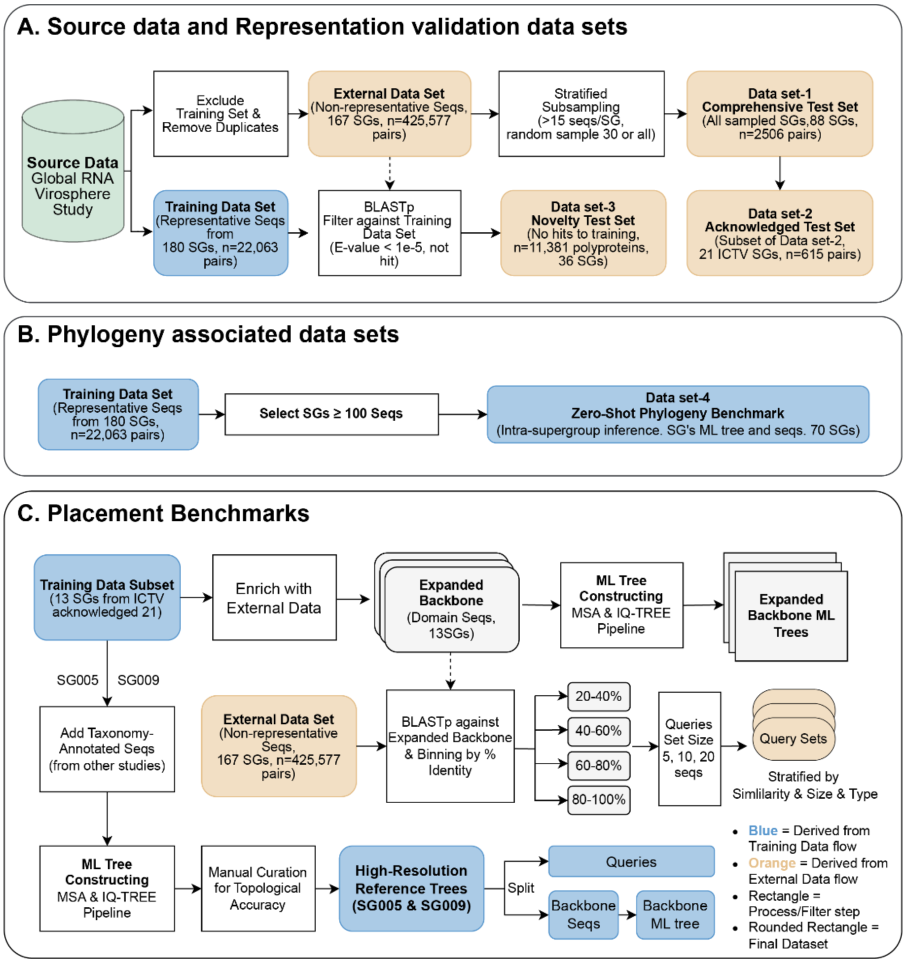


**Fig. S1. Comprehensive data set curation and benchmarking pipeline.**

**(A)** Source data and construction of representation validation data sets (Data sets 1–3). The global RNA virosphere source data was partitioned to ensure strict data set independence, thereby separating the external data set from the training set. Data set-3 isolates highly divergent polyproteins with no detectable sequence similarity to the training set, establishing a rigorous baseline for zero-shot remote homology detection. **(B)** Curation of phylogeny-associated data set (Data sets 4). This data set is designed to evaluate zero-shot topological inference within established viral supergroups. **(C)** Generation of alignment-free placement benchmarks. The upper pathway details the similarity-stratified benchmarking approach, wherein queries from the external data set are binned by percentage identity (20% to 100%) against expanded reference backbones to quantify placement robustness across continuous divergence gradients. The lower pathway illustrates the curation of high-resolution, taxonomy-annotated reference trees (supergroups SG005 and SG009) for direct performance comparison against established alignment-dependent tools.

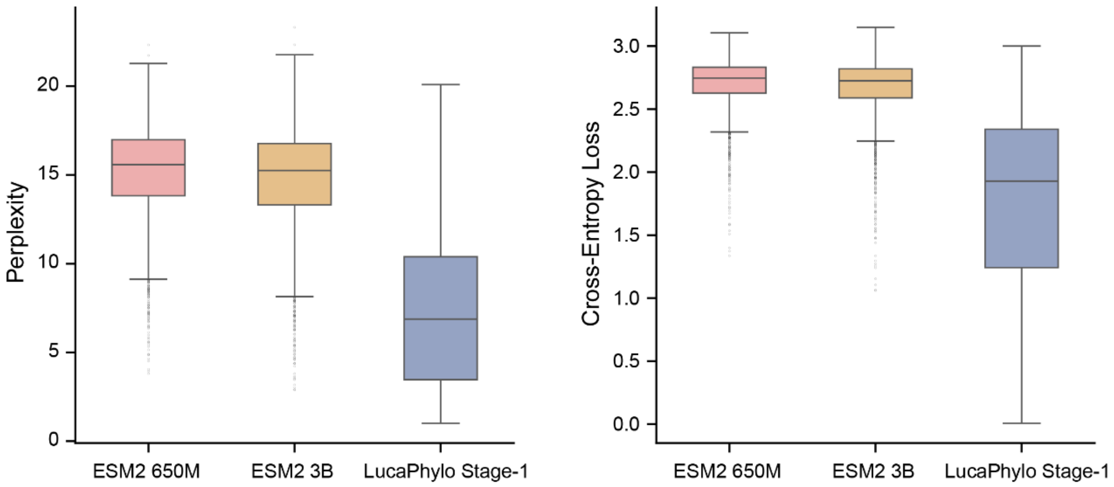


**Fig. S2. Performance evaluation of the Stage-1 model on viral sequences.**

Boxplots comparing the perplexity (left) and cross-entropy loss (right) of general-purpose protein language models (ESM2 650M and ESM2 3B) against the LucaPhylo Stage-1 model. The Stage-1 model was evaluated on Data set-1 and further pre-trained on viral polyprotein sequences. It exhibits substantially lower perplexity and cross-entropy loss with more concentrated variance compared to both baseline models. This indicates its successful adaptation to viral sequence semantics, establishing the stable and optimized initialization fundamentally required for the model's performance in subsequent analytical stages.

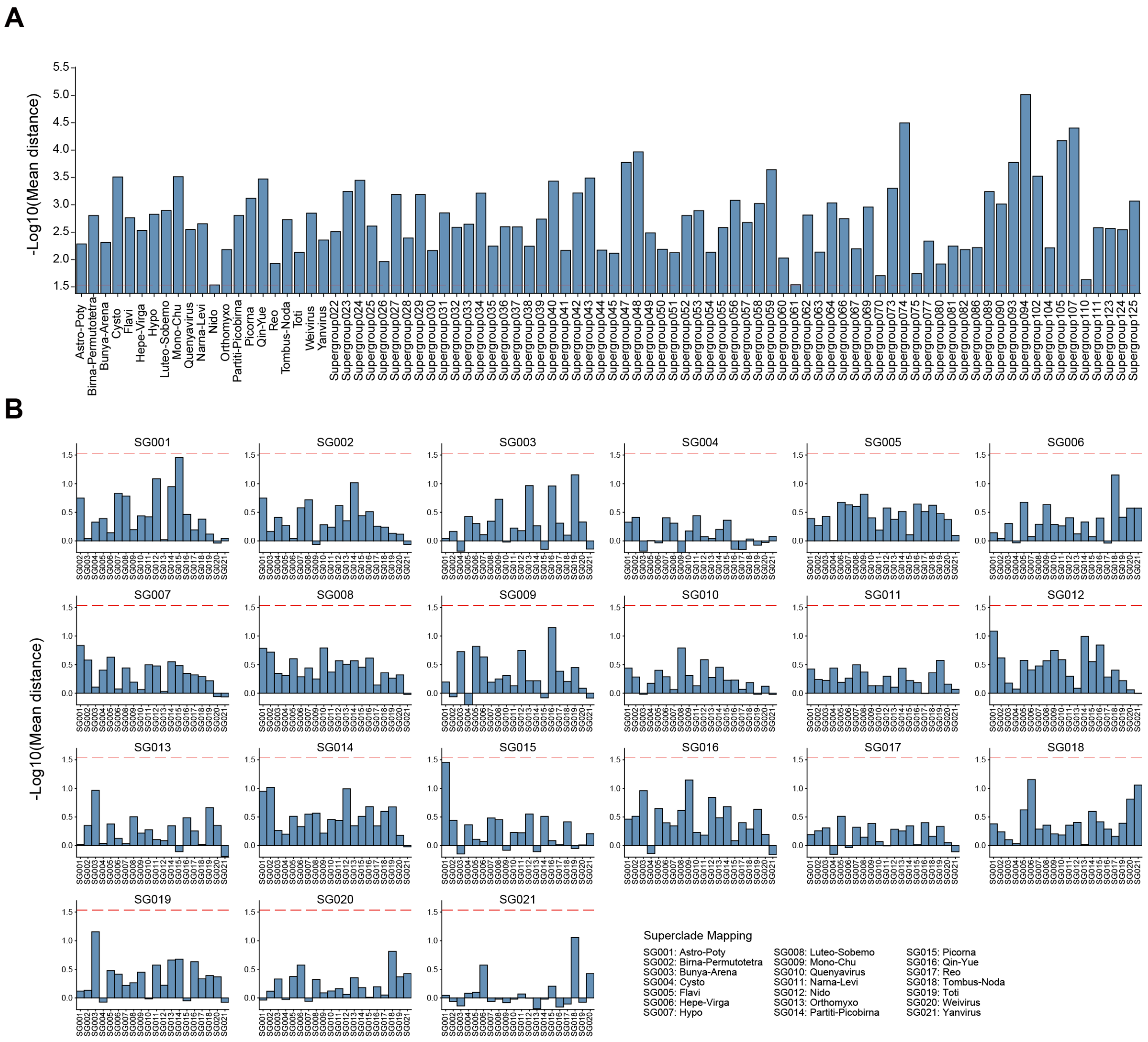


**Fig. S3. Intrinsic separability of viral supergroups in the Stage-2 representation space.**

**(A)** Intra-supergroup similarity scores across 88 viral supergroups evaluated on Data set-1. Scores are logarithmically transformed from pairwise cosine distances among mean-pooled polyprotein embeddings. The red dashed line indicates the global separation threshold, defined as the minimum mean intra-supergroup similarity across all groups, establishing a universal lower bound for within-group sequence cohesion. **(B)** Representative inter-supergroup similarity profiles illustrated using the 21 ICTV-recognized supergroups. Each subplot details the mean similarity of the focal supergroup against all other supergroups in the data set. The red dashed line, projected from **A**, illustrates that inter-supergroup similarities consistently fall below the threshold of intra-supergroup cohesion. Together, these distributions demonstrate that the supergroup-aware contrastive learning autonomously delineates robust geometric boundaries corresponding to broad-scale RNA virus taxonomy.

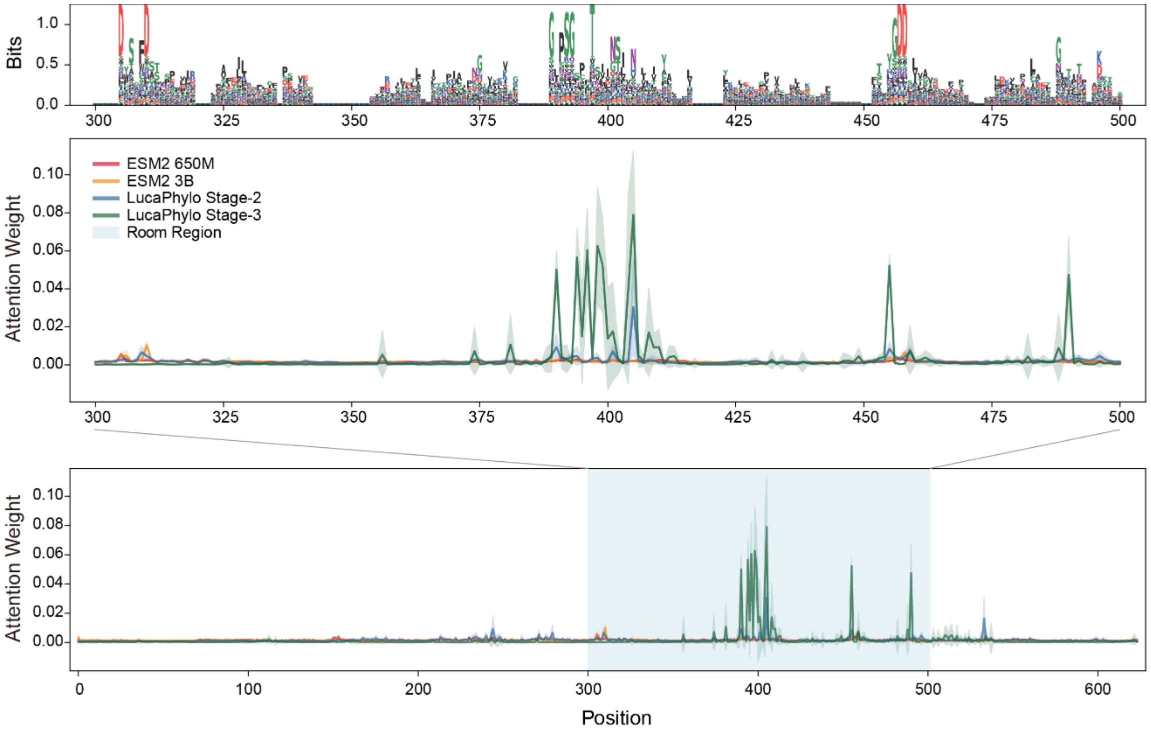


**Fig. S4. Positional consistency between model attention and evolutionarily conserved functional constraints.**

**Top**: Sequence logo derived from the aligned Pfam seed sequences (PF00680), illustrating the sequence conservation spanning the canonical catalytic motifs of the viral RdRP domain. **Middle and Bottom**: Comparative attention profiles across different language models. The bottom panel displays the full-length attention allocation, with the grey shaded area demarcating the RdRP domain. The middle panel provides a magnified view of this specific region. Notably, the LucaPhylo Stage-3 model exhibits distinct attention peaks that are highly concentrated around these key conserved motifs. In addition, these high-attention sites demonstrate strong positional consistency across the protein family. In contrast, baseline general-purpose models (ESM2 650M, ESM2 3B) display diffused and largely uninformative attention patterns across the sequence. This confirms that the cross-modal learning in Stage-3 successfully anchors the model's evolutionary inference to deep functional constraints.

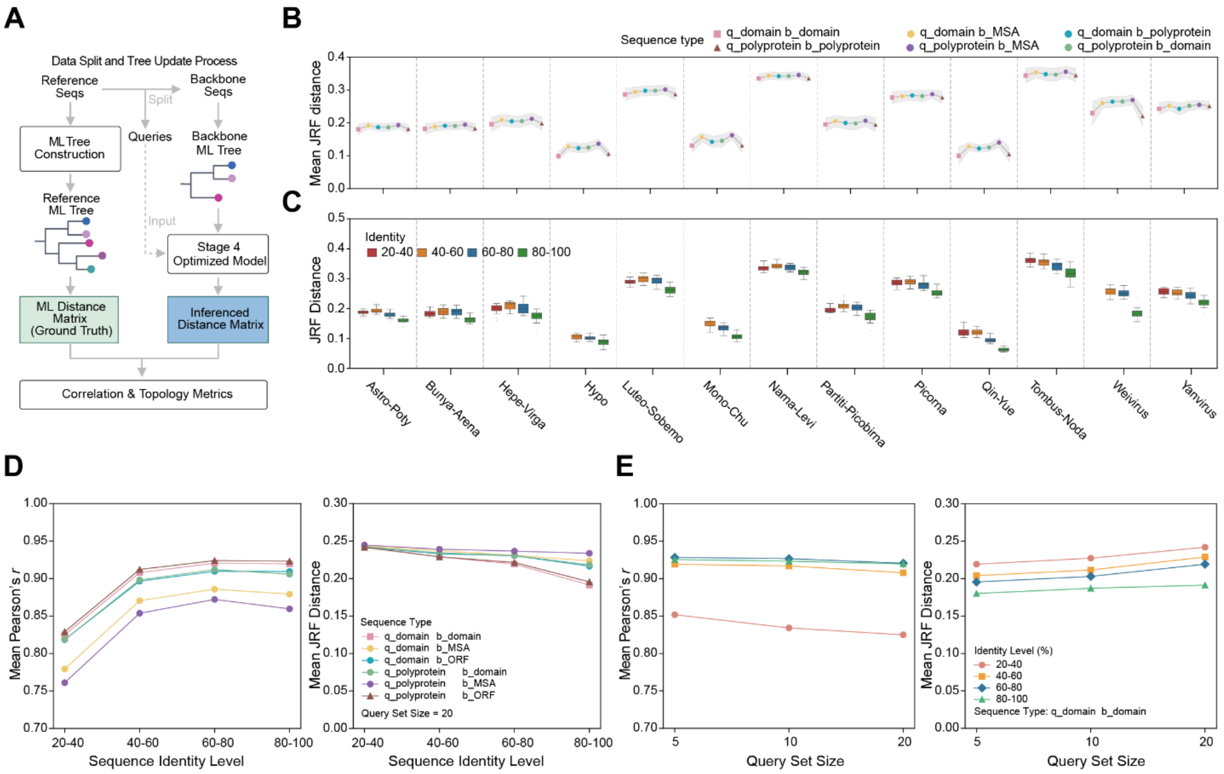


**Fig. S5. Benchmarking of LucaPhylo's alignment-free placement robustness.**

**(A)** Schematic overview of the data split and tree placement process. Reference sequences are partitioned into query and backbone sets to construct ground-truth and backbone ML trees. **(B, C)** Topological divergence of the placement of individual sequences by LucaPhylo with the query set size fixed at 20, complementing the correlation analyses presented in Fig. 4B, C. (B) Mean Jaccard Generalized Robinson-Foulds (JRF) distance across different input modalities, Colors indicate distinct combinations of query (q) and backbone (b) sequence types, with shaded regions representing the standard deviation. Data are aggregated across all sequence similarity levels. **(C)** JRF metric across varying sequence identities, performed using domain sequences for both query and backbone. **(D)** Robustness of alignment-free inference across sequence divergence gradients. Mean Pearson's *r* (left) and mean JRF distance (right) are plotted against binned sequence identity levels for distinct input modalities (query set size = 20). Crucially, the model demonstrates evolutionary fidelity even when both query and backbone inputs consist entirely of unaligned sequences. **(E)** Placement stability assessment across varying query set sizes. Correlation (left) and topological divergence (right) remain highly consistent regardless of the number of simultaneously placed queries (5, 10, or 20 sequences). Analyses were performed using domain sequences for both query and backbone across distinct identity levels.

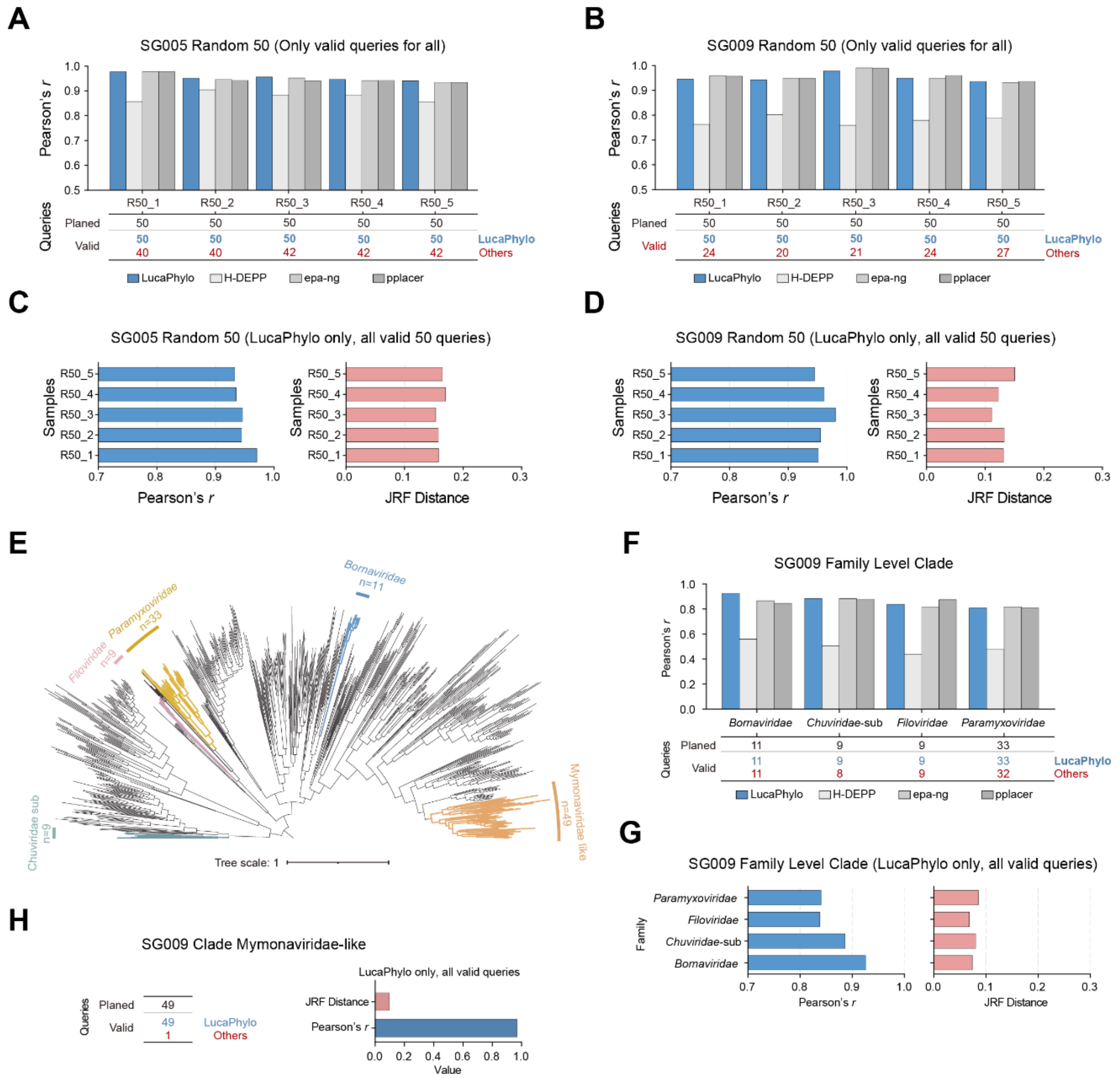


**Fig. S6. Benchmarking alignment-free phylogenetic placement against established alignment-dependent tools.**

**(A, B)** Correlation of inferred evolutionary distances for the placement of individual sequences within the curated supergroup SG005 (A) and SG009 (B) phylogenetic backbones. Performance (Pearson's *r*) is compared between LucaPhylo and three other tools (H-DEPP, EPA-ng, and pplacer) across five independent replicates (50 randomly sampled queries per replicate). The accompanying tables highlight the extensive sequence drop-out experienced by conventional tools due to alignment failures, contrasting with LucaPhylo's 100% sequence retention (valid queries). To ensure a strict and fair comparison, the correlation scores presented for all tools were calculated exclusively using the intersecting subset of queries successfully processed by the conventional tools. (**C, D)** Comprehensive topological divergence and distance correlation metrics for LucaPhylo evaluated on the complete query sets. As conventional tools failed to place all queries, JRF distances—which mathematically require topologically complete tree comparisons—are reported exclusively for LucaPhylo to demonstrate its robustness across the entire, unreduced set of 50 queries per replicate. **(E)** Reference phylogeny of the SG009 (Mono-Chu) supergroup used for clade-placement benchmarks. Colored branches denote the family-level query sets. **(F, G)** Performance evaluations for entire clade placements into the SG009 backbone. Similar to the individual sequence benchmarks, conventional tools suffered sequence drop-out (F); therefore, to maintain fairness, correlation comparisons were rigorously restricted to the overlapping valid queries. In contrast, LucaPhylo successfully processed all sequences, maintaining structural integrity and achieving low JRF distances across the complete and unreduced clades (G). **(H)** Placement performance for the divergent *Mymonaviridae*-like clade. Conventional alignment-based pipelines discarded nearly all sequences (1 valid out of 49 planned). LucaPhylo successfully resolved the placement of this clade, demonstrating high evolutionary correlation and minimal topological divergence.

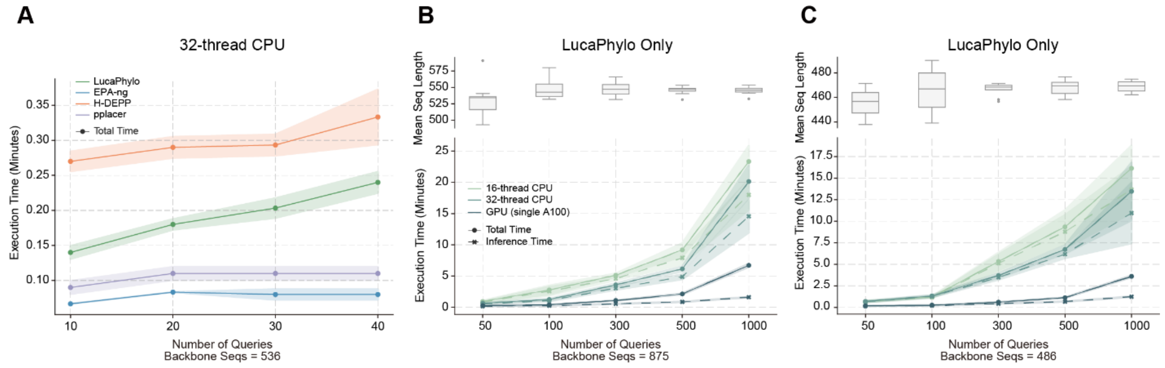


**Fig. S7. Computational efficiency profiling of LucaPhylo for phylogenetic placement.**

**(A)** End-to-end execution time of LucaPhylo compared with baseline alignment-based placement tools (EPA-ng, H-DEPP, and pplacer) on the curated Flavi reference backbone data set described in sequence-level placement (536 sequences). Benchmarks were executed on a standardized 32-thread CPU environment. Solid lines represent the mean execution time, and shaded regions denote the standard deviation across five independent replicates. **(B, C)** Internal computational performance of LucaPhylo across varying query sizes on empirical backbones comprising (B) 875 and (C) 486 sequences. The lower panels display the inference time (dashed lines; encompassing query encoding and distance matrix calculation) and total time (solid lines; incorporating FastME tree reconstruction) across distinct hardware configurations: 16-thread CPU, 32-thread CPU, and a single NVIDIA A100 GPU. Data represent the mean and standard deviation from ten independent replicates. The upper boxplots show the distribution of average sequence lengths for the raw, unaligned queries randomly sampled for each respective cohort size.

#### Legends for Tables S1 to 12

**Table S1**. Metadata for the 180 viral supergroups and detailed sequence composition of the training set.
**Table S2**. Performance metrics for the discrimination of viral supergroups.
**Table S3**. Intra- and inter-supergroup cosine distances on Data set-1.
**Table S4**. Zero-shot inference performance on Data set-4.
**Table S5**. Coefficients of variation for the zero-shot inference performance.
**Table S6**. Ratio of attention weights allocated to the RdRP domain.
**Table S7**. Placement performance across different query and backbone sequence types (query set size = 20, across all sequence identity levels).
**Table S8**. Placement performance across query-backbone similarities and query set sizes (restricted to domain sequences for both query and backbone).
**Table S9**. Benchmark placement performance on individual sequences.
**Table S10**. Runtime comparison for internal phylogenetic placement evaluations.
**Table S11**. Runtime comparison against benchmark methods.
**Table S12**. Benchmark placement performance at the clade level.

#### Legends for Data S1

**Data S1**. Sequence data and metadata for Data set-1 to Data set-4
